# Unexpected plasma gonadal steroid and reproductive hormone levels across the mouse estrous cycle

**DOI:** 10.1101/2023.02.28.530435

**Authors:** Ellen G. Wall, Reena Desai, Zin Khant Aung, Shel Hwa Yeo, David R. Grattan, David J. Handelsman, Allan E. Herbison

**Affiliations:** Department of Physiology Development and Neuroscience, University of Cambridge, United Kingdom; ANZAC Research Institute, University of Sydney, Sydney, Australia; Center for Neuroendocrinology, University of Otago School of Biomedical Sciences, Dunedin, New Zealand

**Author notes:** The authors have nothing to disclose. Grants supporting paper: Welcome Trust (212242/Z/18/Z), Vice Chancellor’s and Newnham College Scholarship, NHMRC Investigator Grant. Author for Correspondence: Allan E. Herbison. Department of Physiology Development and Neuroscience, University of Cambridge, CB2 3EG, United Kingdom.

**Keywords:** Estradiol, Progesterone, Prolactin, Luteinizing Hormone, Estrous Cycle, LCMS

## Abstract

Despite the importance of the mouse in biomedical research, the levels of circulating gonadal steroids across the estrous cycle are not established with any temporal precision. Using liquid chromatography-mass spectrometry, now considered the gold standard for steroid hormone analysis, we aimed to generate a detailed profile of gonadal steroid levels across the estrous cycle of C57BL/6J mice. For reference, luteinizing hormone (LH) and prolactin concentrations were measured in the same samples by sandwich ELISA. Terminal blood samples were collected at 8-hour intervals (10 am, 6 pm, 2 am) throughout the four stages of the estrous cycle. As expected, the LH surge was detected at 6 pm on proestrus with a mean (±SEM) concentration of 11±3 ng/mL and occurred coincident with the peak in progesterone levels (22±4 ng/mL). Surprisingly, estradiol concentrations peaked at 10 am on diestrus (51±8 pg/mL), with levels on proestrus 6 pm reaching only two-thirds of this value (31±5 pg/mL). We also observed a proestrous peak in prolactin concentrations (132.5±17 ng/mL) that occurred earlier than expected at 2 am. Estrone and androstenedione levels were often close to the LOD and showed no consistent changes across the estrous cycle. Testosterone levels were rarely above the LOD (0.01 ng/mL). These observations provide the first detailed assessment of fluctuating gonadal steroid and reproductive hormone levels across the mouse estrous cycle and indicate that species differences exist between mice and other spontaneously ovulating species.

## Introduction

The gonadal steroid hormones have a key regulatory role in controlling reproductive function while also exerting an important modulatory influence upon the activity of almost all other organ systems (1). The hypothalamo-pituitary axis drives marked fluctuations in estrogens and progesterone across the ovarian cycle of mammals. In women, the menstrual cycle typically lasts 28 days and consists of follicular, ovulatory and luteal phases with well-defined stereotypic patterns and levels of circulating reproductive hormones (2). In rodents, the estrous cycle lasts four to five days and can be divided into metestrus, diestrus, proestrus and estrus phases (3). The profiles of reproductive hormones are also clearly documented for the rat estrous cycle (4,5). However, this is not the case for mice where, despite their widespread use in biomedical research, the estrous cycle profiles of circulating estrogens and progesterone are not well established.

The lack of reliable data in the mouse is primarily due to the limitations of steroid hormone immunoassays that include low sensitivity and non-specificity due to cross-reactivity of structurally similar steroid precursors and metabolites. General concerns regarding the specificity of immunoassays for measuring gonadal steroids have been highlighted by academic and clinical communities (6–8) with liquid chromatography-mass spectrometry (LC-MS) now considered to be the gold standard for steroid hormone analysis (9). Nevertheless, prior LC- and gas chromatography (GC)-MS investigations have struggled to detect E2 in mice (10–12). To our knowledge, only a single study has been able to evaluate estrogens and progesterone across the mouse estrous cycle using MS, albeit with GC-MS (13). While valuable, that study examined only a single time point for each day of the cycle.

The present investigation aimed to address the lack of information on circulating gonadal steroid concentrations in cycling female mice by using ultrasensitive LC-MS to measure estrone (E1), 17-β-estradiol (E2), progesterone (P4), testosterone (T) and androstenedione (A4) at 8-hour intervals across the 4-day estrous cycle of C57BL/6J mice. For reference, we also measured luteinizing hormone (LH) and prolactin in the same samples by enzyme-linked immune assay (ELISA).

## Materials & Methods

### Animals

C57BL/6J female mice (>8 weeks old), obtained from Charles River (Margate, UK), were group housed in individually ventilated cages under controlled conditions (12:12 h light/dark cycle, lights on at 05:30 h, 25°C) with environmental enrichment and *ad libitum* access to food (RM1-P, SDS, UK) and water. All procedures were carried out in accordance with the United Kingdom Home Office Animals (Scientific Procedures) Act 1986 (P174441DE) and approved by the Animal Welfare and Ethics Committee of the University of Cambridge.

### Staging of Smears

Vaginal smear samples were collected every morning at approximately 10 am for at least three consecutive cycles to ensure each mouse was actively progressing through the estrous cycle and to determine the cycle stage at death. Mice who displayed irregular cycles were excluded from analysis. The smears were collected by flushing 8 μL of sterile phosphate-buffered saline (PBS) into the vaginal orifice and then collecting the fluid and transferring to a glass slide to air dry before staining with filtered Giemsa (1:1 in Milliq water) and examining under a light microscope. Smear stages were determined by the distribution of either leukocytes, nucleated or cornified cells as reported by (14). As mice can exhibit one or two days of estrus smears, estrus was defined here as mice exhibiting their first estrous smear after proestrus. The vaginal smears were assessed blindly by two independent investigators with 99% agreement. Representative images of estrous, metestrous, diestrous and proestrous smears are shown in (Fig. 1). Estrus was classified by the presence of predominantly cornified cells (Fig. 1A). During metestrus, cornified cells are present and the epithelium starts to become invaded by leukocytes (Fig. 1B). The number of leukocytes becomes more abundant in diestrus (Fig. 1C). Although a reduction is seen in late diestrus alongside a small number of nucleated and cornified cells (Fig. 1D). Proestrus typically consists of a small number of cornified cells with many nucleated epithelial cells present (Fig. 1E).

**Fig. 1.**
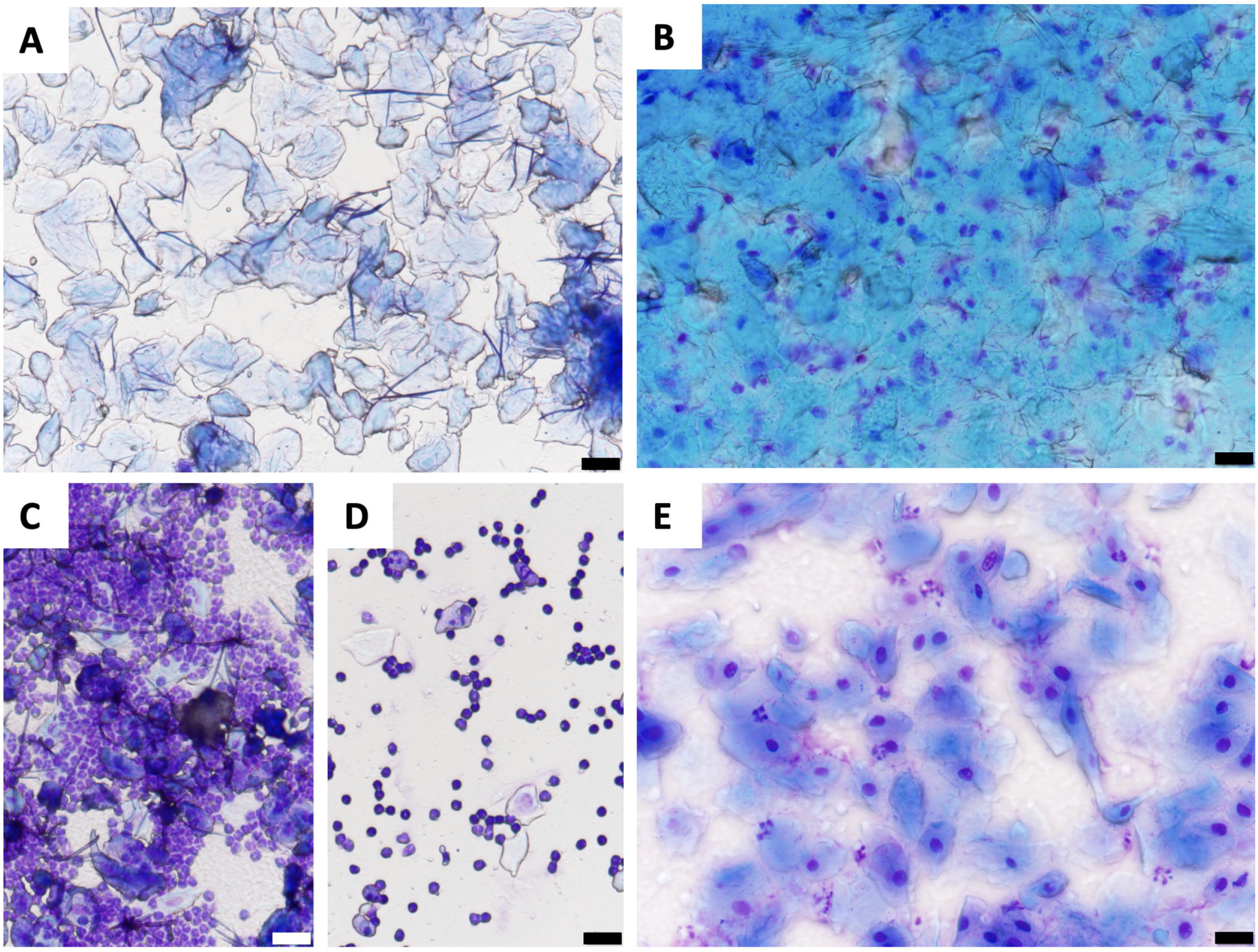
Vaginal cytology of estrous stage smears. A) estrus is represented by cornified epithelial cells, B) metestrus is represented by the presence of cornified epithelial cells and leukocytes, C) early diestrus is represented by a dense appearance of leukocytes, D) late diestrus is represented by a scattered appearance of leukocytes and a small number of nucleated epithelial cells E) proestrus is represented by nucleated epithelial cells and often a small number of cornified epithelial cells Scale bar represents 20 μm.

### Terminal Blood Collection

Terminal blood sampling was performed at 8 hr intervals throughout the four stages of the estrus cycle. Mice ranged between 11 - 19 weeks of age and 16.9 - 25.7 g at the time of terminal blood sampling. The proestrus LH surge occurs approximately 30 mins after lights off (15). To capture this, a 6 pm time point was chosen with the 8 h sampling centered around this at 2 am and 10 am. Immediately prior to terminal bleeding, an additional vaginal smear sample was taken. A lethal dose of pentobarbital (Dolethal, 400 mg/kg, i.p; Vetoquinol UK) was administered and blood was collected within 1 min from the inferior vena cava using a 1 ml syringe flushed with EDTA to prevent coagulation of the blood. Whole blood (approximately 400 μl) was collected and immediately centrifuged at −4°C, RCF 1600g for 5 min. The plasma supernatant was collected, and the pellet was discarded. Plasma for LH ELISA (8 μl) was diluted in (112 μl) 1 x phosphate-buffered saline (PBS) with 0.05% Tween, snap frozen and stored at −4°C. Plasma for prolactin (10 μl) and gonadal steroid (100 μl) was snap-frozen and stored at −80°C prior to analysis by LC-MS at the ANZAC institute. The degree of dilation of the uterine horns was recorded at death.

### Liquid Chromatograph-Mass Spectrometry

The levels of T, A2, E2, E1 and P4 were analyzed using an ultrasensitive LC-MS (16). In brief, hormones were measured in mouse plasma by ultrapressure liquid chromatography-mass spectrometry (LC-MS) in a single batch using a novel, estrogen-selective derivatizing reagent 1,2-dimethylimidazole-5-sulfonyl chloride (DMIS) (23,24). Aliquots (100 μL) of plasma together with standards (estradiol, Cerillant; testosterone, National Measurement Institute (NMI)) and quality control samples were fortified with deuterated d4-estradiol (Cambridge Isotope Lab) and d3-testosterone (NMI) as internal standard, extracted with 1 mL of methyl tert-butyl ether into the organic layer separated from the lower aqueous layer by freezing. Dried extracts were resuspended in sodium bicarbonate buffer and derivatized by addition of DMIS in acetone followed by transfer into a 96-well microtitre plate for 40 μl injection into LC-MS. Chromatography (Kinetex Phenyl Hexyl column, 100 mm × 2.1 mm × 1.7 μm) used a methanol/water gradient with estradiol appearing at 7.07 minutes and testosterone at 5.74 minutes. Eluant was introduced into a mass spectrometer (API 5000 triple-quadrupole mass spectrometer) equipped with an atmospheric pressure photoionization source using negative ionization for estradiol (quantifier transition 271- >145) and positive ionization for testosterone (289->109). For estradiol and testosterone, extraction recoveries were 88%-96% with limits of detection, limits of quantitation and within-run reproducibility (10 replicates) for estradiol were (0.25 pg/mL, 0.5 pg/mL, 5-8%) and for testosterone (10 pg/mL, 25 pg/mL, 4-8%). Spike-recovery of estradiol and testosterone into mouse serum (4 spike levels, 16 replicates) demonstrated high recovery and precision for estradiol (pooled sera 107%, 1.2-4.7%) and for testosterone (111%, 3.4-5.4%).

### EUSA

The sandwich ELISA of Steyn and colleagues (17) was used to measure LH. A standard curve was generated using a serial dilution of mouse LH (reference preparation, AFP5306A; National Institute of Diabetes and Digestive and Kidney Diseases National Hormone and Pituitary Program (NIDDK-NHPP) in PBS-T supplemented with 0.2% BSA. Primary antibody (AFP240580Rb; 1 in 10,000) was purchased from Harbour-UCLS (California, USA) and capture antibody (anti-bovine LH beta monoclonal antibody, 518B7; 1 in 1000) was purchased from UC Davis (California, USA). The secondary antibody (cat#P0448) was purchased from DAKO cytomation (Glostrup, Denmark). The assay sensitivity was 0.03 ng/ml, with intra-assay and inter-assay coefficients of variation of 4.4% and 24% respectively.

Similarly, a previously established sandwich ELISA was used to measure plasma prolactin concentrations (18,19). Reference standard (PRL: 4 μg/ml, AFP6476C, NIDDK-NHPP) was used to generate standard curves ranging from 20 to 0.019 ng/ml by dilution in 0.2% (w/v) bovine serum albumin in PBST. Primary antibody (Rabbit anti-mouse PRL, 1:50,000, AFP131078, NIDDK-NHPP) and capture antibody (Guinea pig anti-rat PRL, 1:2500, AFP65191, NIDDK-NHPP), were purchased from the National Institutes of Health Hormone and Pituitary Program. The secondary antibody was purchased from GE Healthcare Life Sciences (Amersham ECL Rabbit IgG, HRP-linked Ab (from donkey), NA934; used at 1:2000 dilution). The assay sensitivity was 0.04 ng/ml, with intra-assay and inter-assay coefficients of variation of <10%.

### Statistical Analysis

All statistics were performed using (Graphpad Prism 9). Gonadal hormone levels were analysed using a one-way analysis of variance (ANOVA) with a post-hoc Tukey test. The threshold level for statistical significance was set at *P* < 0.05 with data presented as mean ± SEM. Samples below the limit of detection (LOD) were set at LOD/✓2 (20).

## Results

All mice displaying regular 4- or 5-day estrous cycles were included in the analysis except for the proestrus 6 pm time point where only mice exhibiting evidence of an LH surge (LH > 2.0 ng/mL) were used. As proestrus mice exhibit a wide variation in the times at which they initiate the LH surge (15), we only included mice that had commenced the LH surge at 6 pm to maintain consistency around this key time point. All such mice also exhibited greatly dilated uterine horns.

The number of animals (N) for each time point were; Estrus 2am (n=7), 10am (n=4), 6pm (n=5). Metestrus, 2am (n=4), 10am (n=5), 6pm (n=4). Diestrus, 2am (n=9), 10am (n=6), 6pm (n=11). Proestrus, 2am (n=6), 10am (n=8), 6pm (n=8).

### Luteinizing hormone

Mean plasma LH concentrations did not differ significantly between any time points during metestrus, diestrus, and estrus. The mean basal levels for these time points ranged from 0.108-0.755 ng/ml. However, LH levels were significantly augmented during the proestrus 6 pm time point with a mean concentration of 10.66 ± 2.99 ng/ml. These levels were statistically significant compared to all other stages and time points (p<0.0001, ANOVA with post hoc Tukey’s (Fig. 2).

**Fig. 2.**
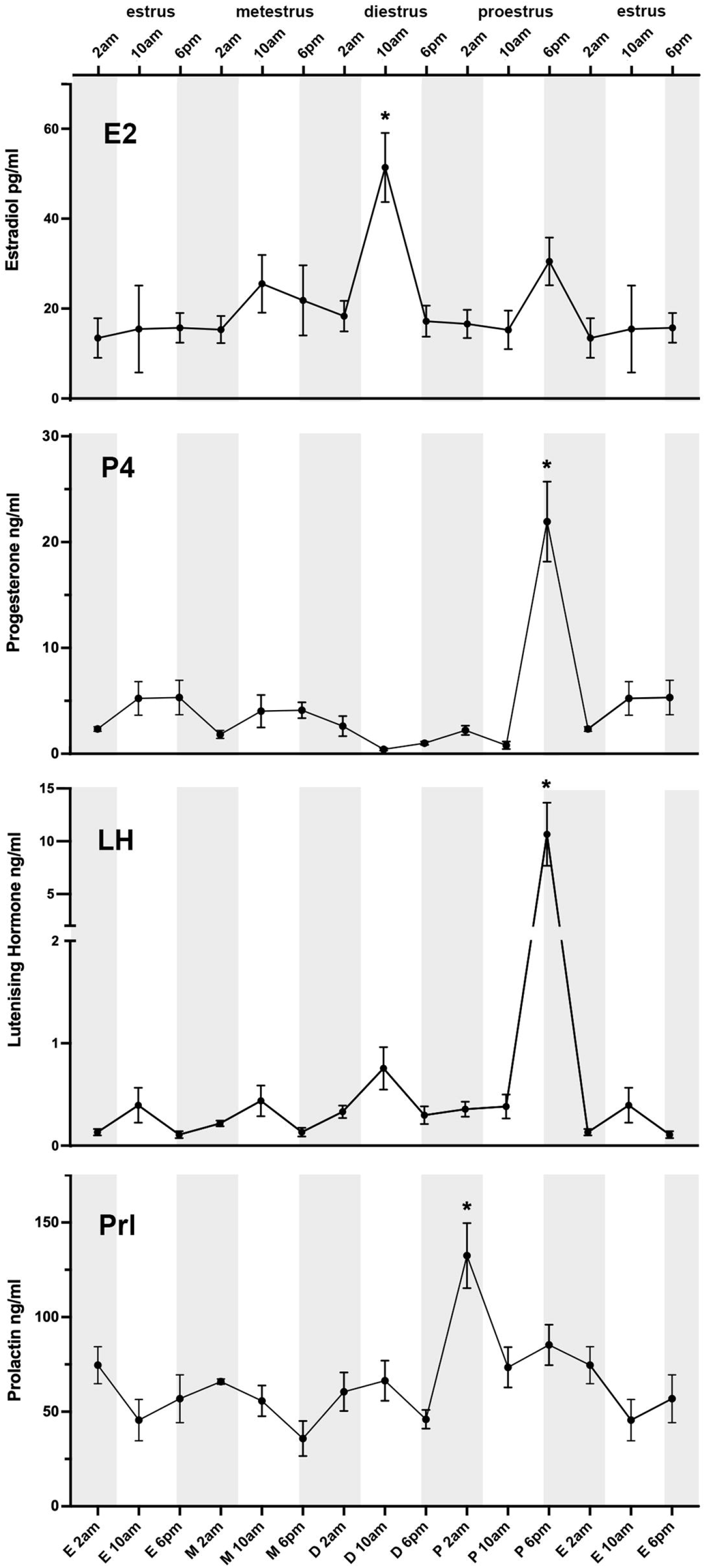
Levels of gonadal and productive hormones across the mouse estrous cycle measured using LCMS. Estradiol (E2) levels on diestrus 10 am are statistically significant compared to all other time points except metestrus 10 am and proestrus 6 pm. Progesterone (P4) on proestrus 6 pm is statistically significant compared to all other time points. Luteinising hormone (LH) on proestrus 6 pm is statistically significant compared to all other data points. Prolactin (Prl) on proestrus 2 am is statistically significant compared to all other time points except proestrus 6 pm. N = 4-11/time point (see text). Grey shaded area represents lights off. Mean ± SEM. ANOVA with post hoc Tukey *p<0.05.

### Prolactin

Mean plasma prolactin concentrations fluctuated around 50 ng/ml and tended to be slightly higher during the day of proestrus compared with earlier days in the cycle, but this did not reach significance. Unexpectedly, however, we observed a large nocturnal peak of prolactin (132 ± 17 ng/ml) at 2.00 am on proestrus. Peak levels at this timepoint were statistically significant compared to all other time points except 6.00pm on proestrus (p<0.05, ANOVA with post hoc Tukey’s (Fig. 2).

### Gonadal steroid hormones

The concentration of P4 peaked at 6 pm proestrus (21.94 ± 3.78 ng/ml) (Fig. 2). These levels were statistically significant to all other time points (p <0.0001, ANOVA with post hoc Tukey’s). A trend towards an increase in P4 was observed at estrus 10 am and 6 pm, metestrus 10 am and 6 pm and diestrus 2 am, however, these were not statistically significant. The lowest levels of P4 were observed at 10 am diestrus (0.42 ± 0.11 ng/ml) (Fig. 2).

Peak E2 levels were detected at 10 am diestrus (51.39 ± 7.67 pg/ml), with E2 levels at 6 pm proestrus being approximately two-thirds this value (30.51 ± 5.30 pg/ml) (Fig. 2). The elevated E2 concentrations observed at 10 am diestrus were significantly different to all other time points except metestrus 10 am and proestrus 10 am (p<0.05 – p<0.001, ANOVA with post hoc Tukey’s) (Fig. 2). The levels of E1 followed a similar pattern to E2 however many values were below the LOD (0.5 pg/ml). Like E2, the concentration of E1 peaked at diestrus 10 am (14.64 ± 4.32 pg/ml) however this showed no significant changes across the estrous cycle (Fig. 3A).

**Fig. 3.**
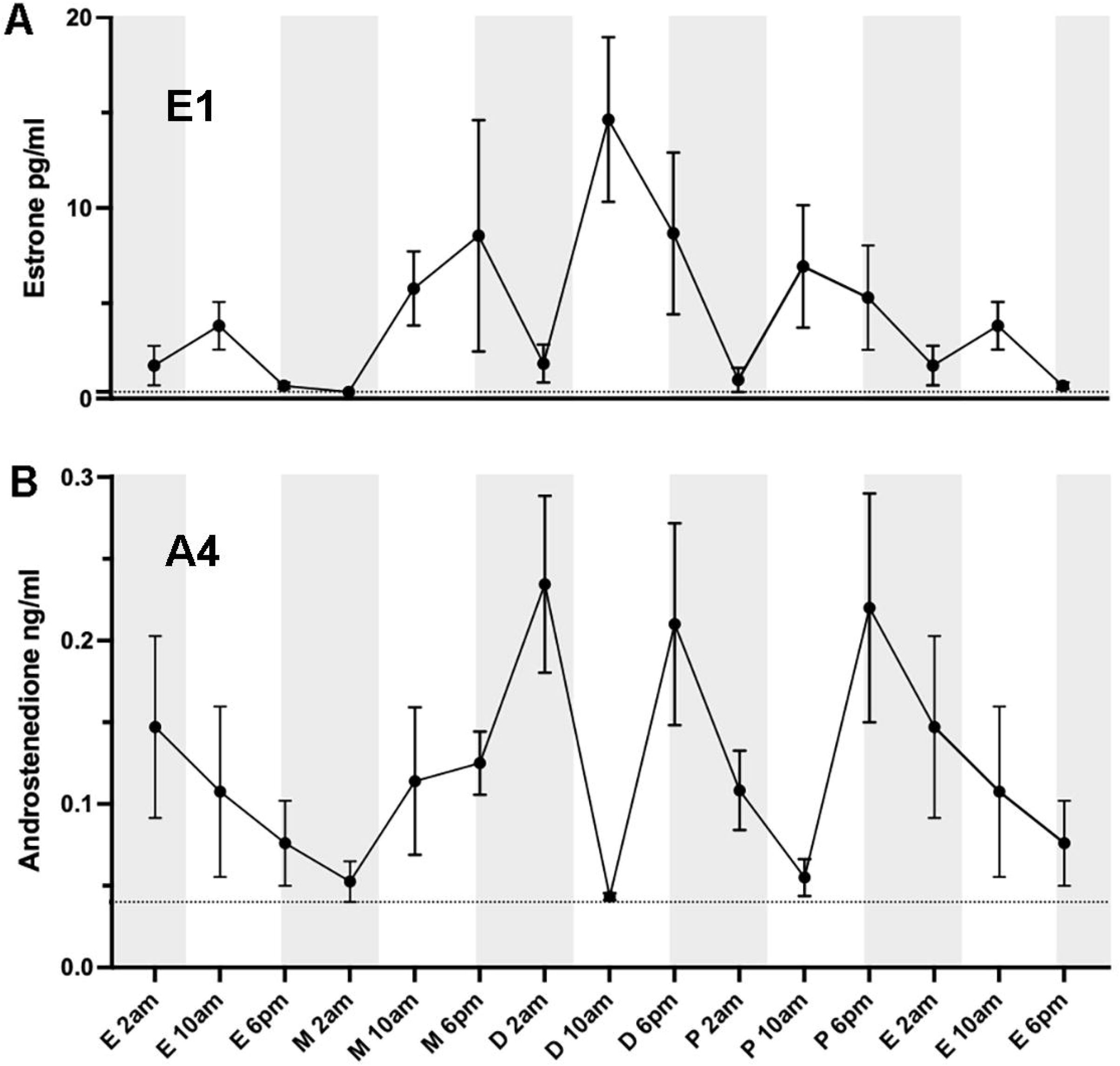
Levels of A) estrone and B) androstenedione across the mouse estrous cycle measured using LCMS. The Grey dotted line represents the LOD/√2. N = 4-11/group. Mean ±SEM.

Virtually all levels of T were below the LOD (0.01 ng/ml). The levels of A4 were often below the LOD (0.05 ng/ml) and showed no significant change across the cycle (Fig. 3B).

## Discussion

These observations provide the first detailed assessment of gonadal steroid hormone levels across the female mouse estrous cycle using LC-MS. Whereas LH and P4 levels peaked at 6pm on proestrus as expected, and in accord with rats (4,5), we found that E2 concentrations were at their highest at 10 am on diestrus and not during proestrus. We also observed an unexpected peak of prolactin concentrations at 2 am on proestrus, 16 hours after the peak of estradiol levels and 16 hours prior to the preovulatory LH surge.

There is wide agreement from human and rat investigations that E2 levels gradually rise across the follicular/diestrous stages to peak on the afternoon or evening of the ovulatory/proestrous phase before falling in the luteal/estrous stage (21). Remarkably, we report here that E2 levels in the C57BL/6J mouse peak on diestrus, the day before ovulation. Using the same LC-MS method, we have previously reported that, as expected, E2 levels are undetectable in ovariectomized C57BL/6 mice (16,22). With respect to the surprisingly low E2 levels on the afternoon of proestrus, we note that all of the 6pm proestrus samples had markedly elevated LH and P4 levels providing proof that these samples were all correctly assigned.

The GC-MS study by Nilsson and colleagues found E2 levels in C57BL/6 mice were the same on proestrus and diestrus but elevated at those times compared to estrus and metestrus (13). A study using commercial immunoassays in C57BL/6 mice reported E2 levels to be at their peak during estrus (23) while a further investigation using Swiss albino mice and an ELISA found maximal E2 levels on proestrus (24). Thus, albeit with different assays, there is substantial variability in reported E2 levels across the mouse cycle. A major caveat for all of these studies is that only a single time point was measured for each day of the cycle. Our present observations showing marked differences between 8-hour time points on the same day indicate the danger of using of single-day measurements.

It is interesting to consider the physiological ramifications of E2 reaching peak concentration on diestrus rather than proestrus in C57BL/6 mice. The rise in estrogens is particularly important in providing estrogen positive feedback to the hypothalamo-pituitary axis to evoke the preovulatory LH surge (25,26). This is considered to represent a classic genomic mode of estrogen action operating through the ESR1 transcription factor (27,28). Accordingly, the surge mechanism requires a long duration E2 signal and high levels do not need to be present at the time of the surge itself (29,30). Intriguingly, the seminal studies of Bronson that established the current E2 treatment regimens used for positive feedback in mice (31,32), employ administration of the high E2 dose at 10 am 32 hours before surge onset. This would be equivalent to the morning of diestrus, the time at which we find E2 levels to peak during the natural estrous cycle.

The preovulatory rise in E2 is also the primary factor driving elevated prolactin secretion during proestrus in the rat (4,5). This action appears to involve a direct action of estradiol on the lactotrophs of the anterior pituitary gland (33). Estrogen-treated ovariectomized rats will express daily prolactin surges in the afternoon, suggesting involvement of an additional circadian timing mechanism stimulating prolactin secretion at that time, driven by releasing factors coming from the hypothalamus (34,35). Previous work has interpreted the absence of an afternoon peak in prolactin in mice to suggest that this second, circadian/hypothalamic contribution to prolactin secretion during the estrous cycle might be lacking in mice, with the prolonged elevation of prolactin seen throughout the day on proestrus likely driven by a pituitary effect of estradiol. The present data may necessitate a reevaluation of this conclusion, as it might be that the timing of this hypothalamic drive to prolactin secretion is earlier in the cycle, driven by the increase of estradiol early on diestrus.

In addition to hormone changes during the estrous cycle, estradiol also stimulates changes in behaviour. There is clear evidence, for example, that estradiol drives an increase in voluntary running wheel activity in rodents (36). It is interesting to note that cyclical changes in running wheel activity during the mouse estrous cycle actually peak during the night preceding proestrus (37) i.e. the night between diestrus and proestrus. This timing is consistent with the peak in estradiol observed in the present study on the morning of diestrus. In contrast, peak running wheel activity in rats occurs in the 24 hours *following* detection of a proestrous smear (effectively in the night between proestrus and estrus) (38). These data too are consistent with the change in running following the peak of estradiol, which is observed in that species in the afternoon of proestrus (4,5).

*We* find that P4 levels dramatically increase at the time of the LH surge and fall dramatically by the early morning of estrus and remain relatively low for the rest of the cycle. As demonstrated in seminal rat studies, the P4 surge occurs at the same time as the LH surge (4,5) and a rise in P4 during proestrus has also been demonstrated in MF1 mice using a radioimmunoassay (39). It is likely that the P4 surge at proestrus 6 pm is driven by the LH surge. Granulosa cells in preovulatory follicles respond to LH and produce P4 in mice (40) and are more sensitive to LH on the morning of proestrus compared to diestrus (41). A single intravenous injection of LH is known to increase concentrations of P4 in rats within 30 minutes (42).

Whereas the proestrous peak in P4 observed here in mice is also observed in rats, we did not observe a significant increase in P4 during estrus/metestrus. There was a non-significant trend towards increased levels of P4 at the 10 am and 6 pm time points in estrus and metestrus. This may represent fundamental species differences in the organization and roles of the luteal phase and estrous. In humans, following ovulation, the ovarian theca and granulosa cells undergo luteinization to differentiate into luteal cells that form the corpus luteum lasting for approximately 2 weeks and become the primary source of P4 in women (43). In contrast, rodents have a very short-lived corpus luteum. They do not exhibit a true luteal phase as P4 levels from the corpus luteum are not sufficient to stimulate decidualization and are rapidly converted to an inactive form 20a-hydroxyprogesterone (44). Rats do however exhibit a second smaller peak in P4 between late metestrus and early diestrus (4,5). Interestingly, only a modest rise in P4 levels during the estrus and metestrus time points was demonstrated in mice. These results may suggest that the P4 increase occurs earlier and is not as pronounced in mice compared to rats (45). Interestingly, mice ovulate earlier in the cycle compared to rats.

We observed that many of the samples had levels of T and A4 that were below the LOD and found no significant variations across the estrous cycle. Low levels of androgens have been reported previously in CB57BL/6 males compared to other strains (46). Additionally, rodents do not synthesize androgens from the adrenal glands, unlike humans, therefore contributing to overall lower plasma levels of androgens (47).

There are several limitations to this study. First, the investigation was undertaken in C57BL/6J mice and may not represent the gonadal steroid levels of other mouse strains. Nevertheless, the C57BL/6 strain is one of the most widely used strains in biomedical research, displaying a high degree of fecundity (48,49) and an understanding of hormone levels in this strain is essential for future work, as well as the re-interpretation of existing studies. Whether other commonly used mouse strains such as 129/SeVe and BALB/c mice have the same gonadal steroid hormone profile will be interesting to assess. While we provide here the most detailed temporal resolution of gonadal steroid levels to date, it may be interesting to examine even finer time points. In particular, the time window from 10 am through to 6 pm may be especially interesting on diestrus and proestrus. Finally, the estrous cycle is classically divided into four stages and vaginal lavage is commonly used to define each stage. However, this implies that each stage fits into, or consistently occurs, within a 24-hour time window which may not always be the case. It is also somewhat variable as to when investigators consider the different stages to occur. In our case we have defined stages beginning and ending at midnight whereas others define the start of each stage in relation to the active and passive phase determined by lighting. Further, mice will often fluctuate between having one- or two-day estrous/diestrus stages during their cycles. Hence, there may be significant subtleties in gonadal steroid hormone profiles associated with these situations that would, again, require more frequent and extensive sampling to reveal.

In conclusion, we provide here a detailed assessment of gonadal steroid hormones levels across the mouse estrous cycle. Surprisingly, we note substantial differences in circulating levels of E2 and prolactin in C57BL/6 mice compared with rats. This study highlights the importance of establishing a detailed hormone profile, which is crucial when using the mouse as a model to study any system affected by gonadal hormones.

